# A wheat/rye polymorphism affects seminal root length and is associated with drought and waterlogging tolerance

**DOI:** 10.1101/463281

**Authors:** Tyson Howell, Jorge I. Moriconi, Xueqiang Zhao, Joshua Hegarty, Tzion Fahima, Guillermo E. Santa-Maria, Jorge Dubcovsky

## Abstract

The introgression of a small segment of wheat chromosome 1BS in the distal region of the rye 1RS arm translocation in wheat (henceforth 1RS^rw^) was previously associated with reduced grain yield, carbon isotope discrimination and stomatal conductance, suggesting reduced access to soil moisture. In this study, we show that the 1RS/1RS^RW^ polymorphism causes differences in root length in field and hydroponic experiments. In the latter, differences in seminal root length were associated with a developmentally regulated arrest of the root apical meristem (RAM). Approximately 10 days after germination, the seminal roots of the 1RS^RW^ plants showed a gradual reduction in elongation rate and stopped growing a week later. Seventeen days after germination, the roots of the 1RS^RW^ plants showed altered gradients of reactive oxygen species and emergence of lateral roots close to the RAM, suggesting a loss of apical dominance. The 1RS/1RS^RW^ isogenic lines also differed in plant biomass and grain yield under normal, terminal drought, and waterlogging field conditions. The differences were larger in fields with reduced or excessive irrigation. These results suggest that this polymorphism may be useful to modulate root architecture and mitigate the negative impacts of excess or reduced water in wheat production.

**HIGHLIGHT:** A wheat/rye polymorphism in chromosome one affects seminal root length and apical dominance and is associated with differences in drought and waterlogging tolerance in the field.

## INTRODUCTION

Approximately 750 million tons of wheat are produced worldwide every year (FAO, 2018), but further increases are required to feed a growing human population. Minimizing yield losses from water stress and waterlogging can contribute to these increases (Mittler and Blumwald, 2010; Tester and Langridge, 2010). One strategy to minimize these losses is the development of crops with root architectures better adapted to different soil types and growth cycles (Manschadi *et al.*, 2006; Voss-Fels *et al.*, 2018). Although some progress has been made in the understanding of root development and architecture in Arabidopsis (Liu *et al.*, 2017; Yu and Luan, 2016), this knowledge is insufficient to engineer useful solutions for specific stress conditions in distantly related grass species (Voss-Fels *et al.*, 2018). One of the traits that contributes to determine plant performance under water limitation is the early capacity of roots to explore the soil at depth (Manschadi *et al.*, 2006); an attribute that could be affected in waterlogged soils (Rich and Watt, 2013).

Rye, a close relative of wheat, is more tolerant to water shortages than wheat, and several studies have pointed to a more robust root system. The translocation of the short arm of rye chromosome one (1RS) to wheat chromosome 1B (1RS.1BL) contributes to above ground biomass (Shearman *et al.*, 2005) and better performance under drought stress (Ehdaie *et al.*, 2012; Ehdaie eř al., 2003; Hoffmann, 2008; Moreno-Sevilla *et al.*, 1995; Zarco-Hernandez *et al.*, 2005). Unfortunately, the 1RS.1BL translocation also brings with it dough stickiness and reduced gluten strength, which negatively affect breadmaking quality limiting its utilization in wheat breeding programs focused on this trait (Fenn *et al.*, 1994). To address this problem a recombinant 1RS chromosome including two wheat 1BS chromosome segment introgressions (henceforth 1RS^ww^) was developed to eliminate the two rye regions generating the problems in bread-making quality (Lukaszewski, 2000).

We introgressed the newly engineered chromosome into the variety ‘Hahn’ using six backcrosses, generating 1RS/1RS^ww^ near isogenic lines (NILs). Previous field trials showed that the Hahn 1RS lines had significantly higher yield and better canopy water status than the 1RS^WW^ NILs in both well-watered and water-stressed environments, although the differences were larger in the latter (Howell *et al.*, 2014). We previously intercrossed the 1RS and 1RS^WW^ lines and generated two additional NILs, one carrying the distal (1RS^RW^) and the other carrying the proximal (1RS^WR^) wheat segment. The two NILs carrying the distal rye region (1RS and 1RS^WR^, henceforth 1RS^xR^) showed significant improvements in grain yield and canopy water status compared to NILs carrying the distal wheat segment (1RS^WW^ and 1RS^RW^, henceforth 1RS^xW^). This indicates that the distal wheat region was the one responsible for the differences. NILs with the distal rye segment also showed higher carbon isotope discrimination and increased stomatal conductance, suggesting that these plants had improved access to the water present in the soil (Howell *et al.*, 2014).

In the winter of 2013, heavy rains caused uneven waterlogging in a UC Davis experimental field that affected the four 1RS NILs at the early tillering stage. Although the affected areas were irregular, the 1RS^xR^ were less affected than the 1RS^xW^ NILs, which showed yellow leaves. Based on this observation, we hypothesized that the 1RS^xR^ lines might have a more extensive root system than the 1RS^xW^ lines, which helped them tolerate both waterlogging in this experiment and water shortages in the previously published experiments (Howell *et al.*, 2014).

In this study, we showed that the wheat-rye polymorphism in the distal region of the 1RS.1BL translocation is responsible for differences in root length in the field. These genotypes also displayed differences in plant biomass (as determined by canopy spectral reflectance) that were generally larger in fields with excessive or reduced irrigation than in fields under normal irrigation. Using hydroponic experiments, we showed that the lines with the distal wheat segment had shorter seminal roots than the lines with the distal rye segment. The shorter roots were associated with an earlier arrest of the root apical meristem (RAM) growth, development of lateral roots closer to the RAM and changes in the distribution of reactive oxygen species.

## MATERIALS AND METHODS

### Plant materials

In this study, we used four near isogenic lines (NILs) that showed differences in grain yield under water stress in a previous work (Howell *et al.*, 2014). The recurrent common wheat parent of these NILs is the cultivar ‘Hahn’ developed by the International Maize and Wheat Improvement Center (CIMMYT). The Hahn cultivar carries the complete 1RS translocation, and the three NILs differed from Hahn either in the presence of a distal interstitial segment of wheat chromatin (henceforth 1RS^RW^), a proximal interstitial segment of wheat chromatin (henceforth 1RS^WR^), or both (henceforth 1RS^WW^) (Howell *et al.*, 2014). The interstitial wheat segments were introgressed from the common wheat cultivar ‘Pavon 76’ to eliminate the *Sec-1* locus from 1RS and to incorporate the *Glu-B3/Gli-B1* locus from 1BS into the 1RS chromosome to improve bread-making quality (Lukaszewski, 2000).

### Waterlogging experiments

Controlled waterlogging experiments were conducted at the University of California field station in Davis, CA (38°32’ N, 121°46’ W) during the 2013-2014 and 2015-2016 growing seasons. An additional experiment was performed in 2014-2015 but it was not analyzed due to severe weed problems. The experiments were planted in November and harvested in June (the harvest year will be used hereafter to designate the experiments).

The two waterlogging experiments were organized in a split-plot randomized complete block (RCBD) design with four blocks in 2014 and three blocks in 2016. Within each block, the main factor was irrigation treatment, and within each irrigation treatment – block combination, the Hahn1RS, 1RS^WW^, 1RS^RW^, and 1RS^WR^ genotypes were used as sub-plots. The average of 1RS and 1RS^WR^ was compared with the average of 1RS^WW^ and 1RS^RW^ to determine the effect of the distal rye and wheat chromosome segments.

In the 2014 field experiment, each block included two different irrigation regimes as main plots. The first one was based on plant needs and normal practices in California Sacramento Valley and the second one consisted of artificial flooding twice a week starting in late January and ending in late March 2014 during the tillering stage. Water was applied via flood irrigation, and the schedule was set so that the soil profile remained saturated. While plants were not kept fully or partially submerged, there were persistent pools of water on the soil surface indicating a waterlogged environment. Each genotype was planted in three adjacent 1 m rows (experimental unit) at a rate of 30 grains per row. These experimental units were replicated six times within each of the four blocks in an RCBD pattern and were used as sub-samples. At the end of the season, each set of three rows was harvested and grain yield was recorded in grams. The average of the six subsamples was used as a single data point in the statistical analysis. Canopy Spectral Reflectance (CSR) measurements were taken for all subsamples on two days (4/17and 4/30). Subsamples were averaged within days, and day averages were used as repeated measures.

Canopy spectral reflectance (CSR) measurements were taken with the “ASD HandHeld 2 Pro” spectrometer from Malvern Panalytical. Measurements were taken using a “scanning” method in which 50 measurements were taken on a single plot and averaged together to give a single representative reflectance spectrum. From these measurements, differences in biomass between genotypes were estimated using the Normalized Difference Vegetation Index (NDVI), which was calculated using the formula (R900 nm-R680 nm)/(R900 nm+R680 nm).

In the 2016 field experiment, each block included three irrigation treatments. The first one was a normal irrigation with water provided as needed. The waterlogging treatment included flood irrigations three times a week that were applied as described above. Irrigations were initiated at the beginning of February (later than in 2014 due to a wet winter) and ended at the end of February. In the drought treatment, irrigation ceased in late March before the booting stage. Within each block – treatment combination, each genotype was machine sown at a density of 3 million grains per hectare in 2.23 m^2^ plots (1.83 x 1.22 m), which were combine-harvested at maturity. In 2016, CSR measurements were taken as described above on four different days (3/24, 4/6, 4/13, and 4/28). Days were used as repeated measurements and were analyzed as sub-sub-plots in an RCBD split-split plot design using conservative degrees of freedom for days and all their interactions (this was not necessary in 2014 because there were only 2 days and a single degree of freedom). After the CSR measurements were completed, an irrigation pipe ruptured flooding several sections of the experiment in an irregular pattern (4/29) and increased variability in the final yield measurements.

### Root depth experiments

The field experiment to estimate root length was conducted at the University of California field station in Davis, CA, which has deep Yolo loam soils (fine-silty, mixed, superactive, nonacid, thermic Mollic Xerofluvent). The field was organized as a randomized complete block design with six blocks and four genotypes per block. Plots were machine sown at a density of 3 million grains per hectare in 4.5 m^2^ plots in November 2016.

To obtain soil samples at specific depths and avoid differential soil compaction, we excavated ~2 m deep trenches cutting perpendicular across the middle of plots and including complete blocks one (3/3/2017, North side), three (3/20/2017, North side) and six (3/9/2017, South side) to expose the root system. We then took horizontal soil core samples from the center of each block at 20 cm intervals using a thin-walled copper pipe (5.08 cm diameter x 35 cm long= 709.4 cm^3^). Core samples were taken from 20 to 140 cm in the first block and from 20 to 180 cm in blocks three and six (Supplementary Fig. S1) after we discovered the presence of roots at 140 cm in block 1. Plants were at the tillering stage at the time of the root sampling.

### Determination of root parameters in soil samples

Soil core samples were washed using a hydro-pneumatic elutriation system from Gillison’s Variety Fabrications, Inc. (Smucker, McBurney, and Srivastava 1982). After washing and sorting roots from other organic matter, we suspended the roots in water and scanned them using an EPSON Expression 11000XL flatbed scanner. Scanned root images were analyzed using the WinRhizo software package. We also attempted to measure root biomass, but the small biomass of the dried roots combined with small stray soil contaminants and changes in ambient moisture made it difficult to quantify root biomass, so only the WinRhizo measurements are reported. The 20 cm sampling point was not used because the large amount of root biomass and organic matter present in these samples made them difficult to clean and measure accurately.

Since all root measurements were performed using soil cores of identical volume (709.4 cm^3^) we will refer to these measures as densities. Differences in total root length, surface and volume density, average root diameter, and root tips and fork densities were analyzed using a split plot design with genotypes (average of 1RS and 1RS^WR^ vs. average of 1RS^WW^ and 1RS^RW^) as main plots and depth as subplot. This is a very conservative statistical analysis because it reduces the df for genotype from 3 to 1. Therefore, we also compared the two same pairs of genotypes using statistical contrasts in an ANOVA including all four genotypes. To account for the inability to statistically randomize depths, we used a conservative estimate of the degrees of freedom (df) for subplots and for the interaction between subplot and main plot. Conservative df were calculated by dividing their df by the number of subplots. This strategy is similar to the one use for repeated measures in time and does not affect comparisons among main plots (genotypes), which are the main objective of this study. Homogeneity of variance and normality of the residuals was confirmed for all the individual ANOVAs performed at each depth for all parameters. When necessary, data was transformed using power transformations to satisfy ANOVA assumptions.

### Hydroponics experiments

The hydroponic experiments were performed in growth chambers set at 22-23 °C with a photoperiod of 16 h light / 8 h dark provided by a fluorescent light source supplemented with incandescent lighting. In all experiments, grains were imbibed at 4°C for four days, after which they were placed at room temperature. Once the majority of the grains had germinated and the coleoptiles had emerged, healthy germinated seedlings were transferred to a mesh suspended on water (UCD) or CaCl_2_ (0.5 mM, Argentina). Four days later, healthy seedlings were transferred to tanks with growth solutions (KH_2_PO_4_ 0.2 mM, MgSO_4_.7H_2_O 1.0 mM, CaCl_2_1.5 mM, KCl 1.5 mM, H_3_BO_3_ 1 μM, (NH_4_)_6_Mo_7_O_24_.4H_2_O 0.05 μM, CuSO_4_.5H_2_O 0.5 μM, ZnSO_4_.7H_2_O 1 μM, MnSO_4_.H_2_O 1 μM, FeEDTA 0.1 mM, Ca(NO_3_)_2_ 1.0 mM, (NH_2_)_2_SO_4_ 1.0 mM). After removing the remaining grain, seedlings were wrapped at the crown with foam and inserted in holes precut in a foam core board placed on top of the solution. Nutrient solution was changed two to three times a week for the duration of the experiment. Particular details of the methods used in the two laboratories are described below:

#### Davis, CA, USA

The experiments were performed in 13 L hydroponic tanks (ULINE part number S-19507) containing the nutrient solution. Twenty-four seedlings were placed in each hydroponic tank in a six by four pattern. All genotypes were included in one tank, and if necessary, multiple tanks were used as replications.

In the experiments aimed to study the effect of different sources of nitrogen and of their external concentrations, seedlings were grown in normal growth solution for seven days (from 5 to 12 DAG) and then transferred to low nitrogen solution for another 10 days (12 to 22 DAG). Roots were measured and imaged with a flatbed scanner at 22 DAG.

#### Chascomús, Buenos Aires Argentina

The CaCl_2_ (0.5 mM) from the tank where the grains were germinated was replaced by nutrient solution on the 4^th^ day. On the 5^th^ day, plants were transferred to 0.350 L pots containing aerated nutrient solution, with each pot (plant) being a replicate. Pots were rotated every two days to ensure that they occupied different positions within the growth chamber. For the root time course, the length of the second longest seminal root was measured four hours after the start of the light period, starting 6 DAG.

### Nitro blue tetrazolium (NBT) and 2’,7’-dichlorofluorescin diacetate (DCF-DA) staining

Five centimeter root sections were carefully excised from the second longest root of 1RS and 1RS^RW^ plants 17 DAG. For the NBT staining this segment was placed for 90 min in a 0.1 mg/mL NBT solution dissolved in 200 mM potassium phosphate buffer, pH 7.6, in darkness. For the DCF-DA staining root segments were placed in the same buffer supplemented with 10 μM DCF-DA for 60 min, in darkness. Roots segments stained with NBT (detecting superoxide anions) or DCF-DA (detecting hydrogen peroxide, peroxynitrite and hydroxyl radicals) were washed in the same buffer for 30 min and placed on a slide. Roots were observed using a Zeiss Discovery.V20 (Carl Zeiss MicroImaging, Germany) stereomicroscope equipped with a coaxial fluorescence mechanism. Pictures were obtained with an Axiocam 512 color (Carl Zeiss MicroImaging, Germany). The images were processed with the ImageJ software obtaining a longitudinal profile of color and fluorescence intensities.

## RESULTS

### 1RS^xW^ lines showed less tolerance to waterlogging than 1RS^xR^ lines in the field

To test if the 1RS^xR^ lines have better tolerance to waterlogging than the 1RS^xW^ lines we performed controlled waterlogging experiments in 2014 and 2016 using the four Hahn NILs. The analysis of the data using the four different genotypes showed that the NDVI and yield responses of the lines carrying the distal rye chromosome segment (1RS + 1RS^WR^ =1RS^xR^) were similar to each other and different from the lines carrying the distal wheat chromosome segment (1RS^WW^ + 1RS^RW^ =1RS^xW^), which were also similar to each other (Supplementary Fig. S2). These results were consistent across years and irrigation treatments, so for simplicity we grouped the genotypes with the same distal segment for the following statistical analyses and figures.

#### 2014 field experiment

The repeated measures ANOVA analysis (two days) of the RCBD split plot experiment harvested in 2014 showed significant differences between control and waterlogging conditions (main treatment) for NDVI *(P =* 0.0025, Supplementary Table S1), but the effect of the treatment on yield was not significant (P = 0.1556, Supplementary Table S3). For both traits we detected highly significant differences between genotypes (sub-plots, *P* <0.0001), with the 1RS^xR^ lines showing higher NDVI and grain yield than the 1RS^xW^ lines (Fig. 1). The NDVI differences between genotypes under waterlogging conditions were larger than under normal irrigation. This was reflected in a significant interaction between waterlogging and genotype (P = 0.0034, Supplementary Table S1) that can be visualized in the interaction graph as lack of parallelism between lines (Fig. 1). The same trend was observed for yield (Fig. 1), but the interaction was not significant (P =0.1101, Supplementary Table S3), likely due to the conservative statistical analyses used for these tests (see Material and Methods) and the higher variability of the yield data.

**Fig. 1.**
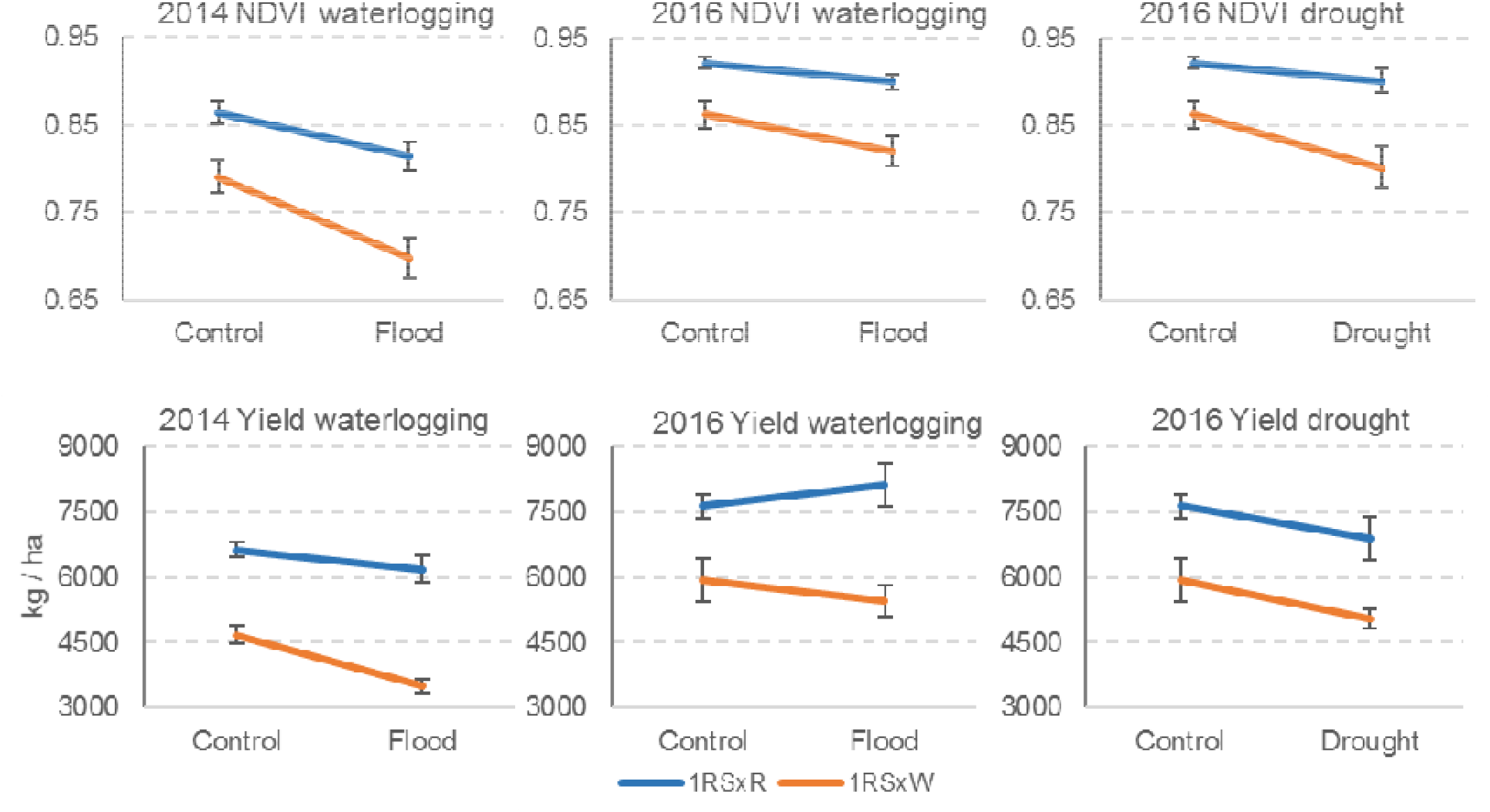
Plant biomass estimated by NDVI (top row) and yield (bottom row) under normal irrigation (control), waterlogging and drought conditions. Lines with the distal rye segment (1RS^xR^) showed significantly higher NDVI and yield values (P<0.0001) than the lines with the distal wheat segment (1RS^xW^) in all experiments. Differences between genotypes were generally larger under water stress conditions. Error bars are SE of the means across blocks.

Since the Genotype x Day interaction for NDVI was significant, we also performed separate statistical analyses for each of the two days. Each of the two days show similar results, with significant differences between treatments, genotypes and genotype x treatment interactions (Supplementary Table S2).

#### 2016 field experiment

This RCBD split-split-plot experiment included three irrigation treatments (control, waterlogging, and drought). To facilitate the comparisons with the 2104 experiment and to simplify the statistical analysis of the interactions, we present the ANOVA tables for waterlogging (Supplementary Tables S4-6) and drought separately (Supplementary Tables S7-9), although they share the same control treatment.

In the waterlogging experiment, we observed significant differences between treatments for NDVI (P = 0.0010, Supplementary Table S4) but not for yield (P = 0.9772, Supplementary Table S6). The unanticipated rupture of an irrigation pipeline (after all NDVI measures were completed) affected the treatments and blocks in irregular patterns leading to increased variability of the grain yield results. This increased variability might have contributed to the lack of significant differences in grain yield between treatments. In spite of this incident, the differences between genotypes were highly significant for both NDVI (P<0.0001, Supplementary Table S4) and grain yield (P =0.0054, Supplementary Table S6). Similar to the 2014 experiment, the 1RS^xR^ lines showed higher NDVI and grain yield than the 1RS^xW^ lines (Fig. 1). We observed a consistent trend for larger NDVI differences between genotypes in the waterlogging treatment than in the control (Fig. 1), but the interaction between waterlogging and genotype was marginally non-significant in the overall NDVI analysis *(P =* 0.0783, Supplementary Table S4) and not significant for yield (P = 0.2888, Supplementary Table S6).

For NDVI, the individual ANOVAs by day showed marginally non-significant interactions for the measurements taken in the middle of the waterlogging experiment (April 6 *P =* 0.06 and April 16 *P =* 0.07), but not for those closer to the beginning and end of the measurement window (March 24 *P =* 0.91 and April 28 *P =* 0.16, Supplementary Table S5). The differences between genotypes were highly significant for all four individual ANOVAs (Supplementary Table S5).

No significant differences were detected between the drought and control treatments for NDVI (P = 0.1130) and grain yield (P = 0.4587). By contrast, highly significant differences were detected between genotypes for both parameters (NDVI *P* = 0.0002, Supplementary Table S7, grain yield *P* = 0.0088, Supplementary Table S9). We also detected a significant treatment by genotype interaction for NDVI (P = 0.0308, Supplementary Table S7) but not for yield (P = 0.8561, Supplementary Table S9), likely due to the additional variability generated by the irrigation pipe incident. In the four individual ANOVAs by day, the 1RS^xR^ lines showed higher NDVI and grain yield than the 1RS^xW^ lines (Fig. 1, Supplementary Fig. S2, Supplementary Table S8).

Taken together, these results confirmed the better performance of the 1RS^xR^ lines compared to 1RS^xW^ lines, under control, waterlogging and water deficit conditions. The differences between genotypes tended to be larger under excessive or insufficient irrigation than under control irrigation (Fig. 1), which suggests that the presence of the distal 1RS segment helped to mitigate the negative impacts of waterlogging and drought on biomass accumulation and grain yield.

### 1RS^xW^ lines have shorter roots than 1RS^xR^ lines in the field

#### Total root length density

The graph for total root length density (total root length in a 709.4 cm^3^ soil volume) at different soil depths showed consistent differences between genotypes grouped by the distal rye or wheat segments (Fig. 2A). The total root length densities were consistently higher in the 1RS^xR^ than in the 1RS^xW^ NILs through the soil profile, with the largest absolute differences detected at 40 cm (Fig. 2A).

**Fig. 2.**
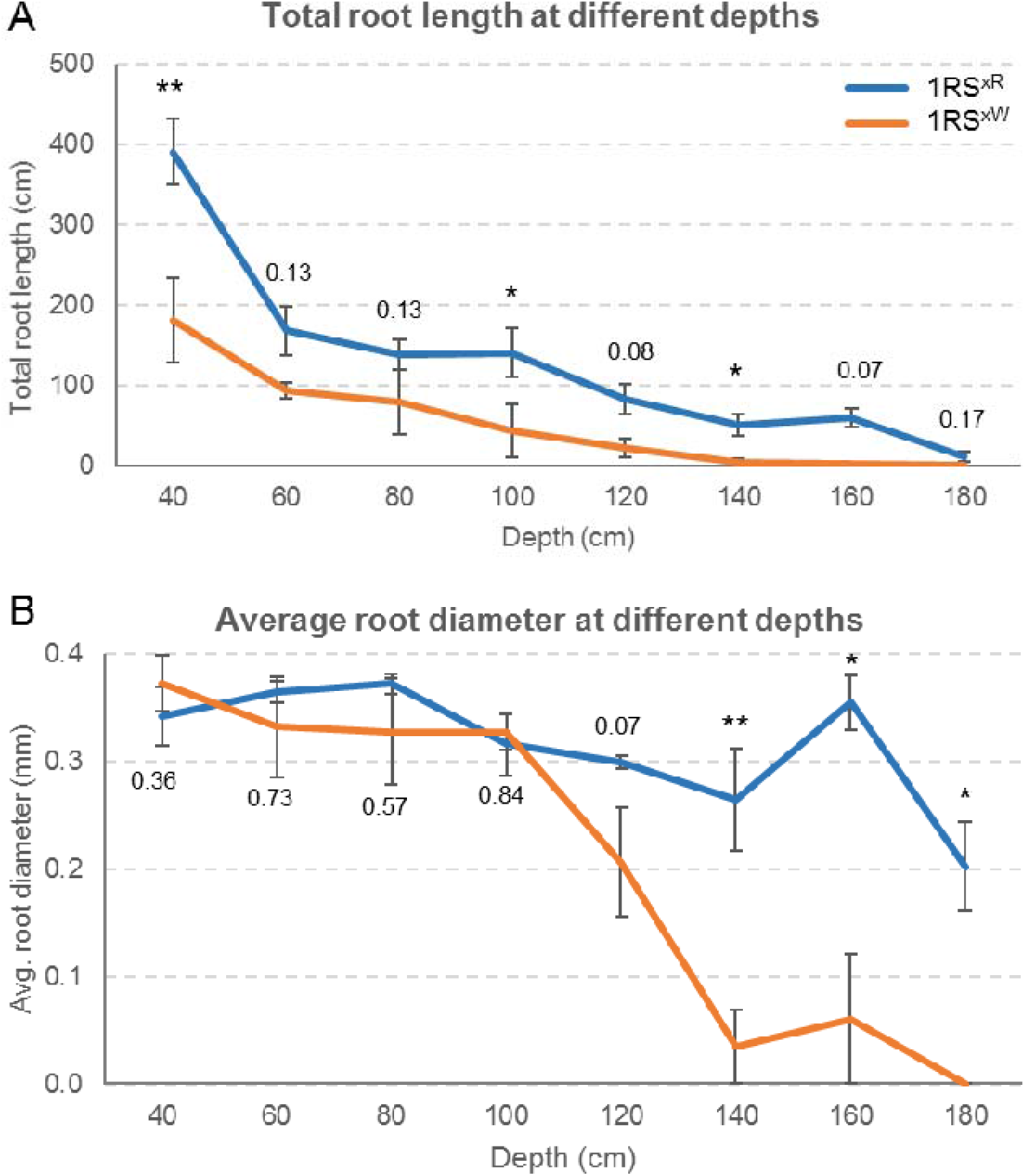
Decrease in total root length density and average root diameter in 1RS^xW^ and 1RS^xR^. **A)** Total root length density. Relative changes are presented in Supplementary Fig. S3A and S3B. **B)** Average root diameter. *P* values represent differences between 1RS^xW^ and 1RS^xR^ at individual depth (Tables S12 and S15). Error bars represent SE of the means across blocks. * = *P* <0.05, ** = *P*<0.01, *** = *P*<0.001.

The overall ANOVA for total root length density showed significant differences between the averages of the 1RS^xR^ and 1RS^xW^ genotypes (P =0.0263, Supplementary Table S10). The limited degrees of freedom resulting from the averaged genotypes make this analysis very conservative. To increase the power of the analysis, we performed a statistical contrast between the two lines with the distal rye segment and the two lines with the distal wheat segment in a separate ANOVA including all four genotypes. In this analysis with a higher number of degrees of freedom, the difference between the 1RS^xR^ and 1RS^xW^ genotypes was highly significant (P = 0.004, Supplementary Table S11).

The overall ANOVA showed a marginally significant interaction between genotype and depth (*P* = 0.0497), but it become not significant when adjusted using conservative degrees of freedom (*P* = 0.190, Supplementary Table S10). The total root length density decreases with depth, which minimizes the absolute value of the differences between genotypes at more extreme depths. To address this limitation, we explored the differences between genotypes expressed as a percent of the 1RS^xR^ values (((1RS^xR^ – 1RS^xW^) / 1RS^xR^) *100). This analysis showed that the relative difference in total root length between 1RS^xW^ and 1RS^xR^ plants was larger at the deepest sampling points (Supplementary Fig. S3A, regression between depth and percent difference *P* = 0.002). No roots were detected for 1RS^xW^ at 180 cm.

Finally, we performed individual ANOVAs by depth and detected significant differences between genotypes at three depths and marginally non-significant differences at two (Fig. 2A, Supplementary Table S12). In summary, Fig. 2A and the statistical analyses in Supplementary Tables S10 to S12 indicated that the 1RS^xR^ lines have more extended and deeper root systems than the 1RS^xW^ lines.

#### Average root diameter and combined root traits

In the overall split-plot ANOVA for average root diameter, the effect of depth was significant, even when using conservative degrees of freedom (*P*= 0.02, Supplementary Table S13). The difference between the average 1RS^xR^ and 1RS^xW^ genotypes was marginally non-significant in the conservative analysis (*P* = 0.0649, Supplementary Table S13), but was highly significant in the statistical contrast between the pairs of genotypes in the ANOVA including the four genotypes (*P* = 0.006, Supplementary Table S14). In the analyses by individual depths, we observed similar average root diameters for the 1RS^xR^ and 1RS^xW^ genotypes between 40 and 100 cm, but those values started to differentiate at 120 cm (*P*= 0.07) and were significant at 140, 160 and 180 cm. At all these depths, the average root diameter was larger in the 1RS^xR^ than in the 1RS^xW^ lines (Fig. 2B, Supplementary Table S15).

A similar result was observed also when the differences in root diameter were calculated as the decrease in 1RS^xW^ relative to 1RS^xR^. In this graph, a sharp increase in the effect of genotype was observed below the 100 cm depth (Supplementary Fig. S3B, regression between depth and relative root diameter *P* = 0.002). Taken together, these results suggest that the main seminal and/or adventitious roots from the 1RS^xR^ lines reach deeper in the soil than the1RS^xW^ lines, for which only thinner roots are found deeper in the soil profile.

The other root traits measured with WinRhizo showed a profile similar to the one observed for total root length density (Supplementary Fig. S4). This is not surprising because root surface and volume density are a function of root length and diameter, and the number of root tips and forks is mainly affected by the abundance of roots. The contrasts between 1RS^xW^ and 1RS^xR^ in the ANOVAs using the four genotypes were significant for root surface (P =0.005), root volume (P =0.008), number of tips (P =0.02) and number of forks densities (P =0.003, Supplementary Fig. S4). Significant differences were also detected in three to four of the ANOVAs for individual depths in each of the traits (Supplementary Fig. S4).

### 1RS^xW^ lines have shorter primary roots than 1RS^xR^ in hydroponic cultures

The changes in total root length density and diameter with soil depth include both seminal and adventitious roots. To see if the differences observed between the 1RS^xW^ and 1RS^xR^ lines in the field could be also detected at early growth stages, when the root system is dominated by seminal roots, we performed hydroponics experiments.

#### Differences in root length

In the hydroponic experiments, the 1RS and 1RS^WR^ lines showed significantly longer seminal roots than the 1RS^RW^ and 1RS^WW^ lines (Fig. 3A). Since root length was not significantly different between lines with the same distal segment, those lines were averaged for the statistical analyses (Fig. 3B). The differences in seminal root length between the 1RS^xW^ and 1RS^xR^ plants were detected in experiments using two sources of nitrogen (nitrate and ammonium) at two different concentrations (2.0 and 0.2 mM, Fig. 3B), which indicates that these differences are robust across different ionic environments.

**Fig. 3.**
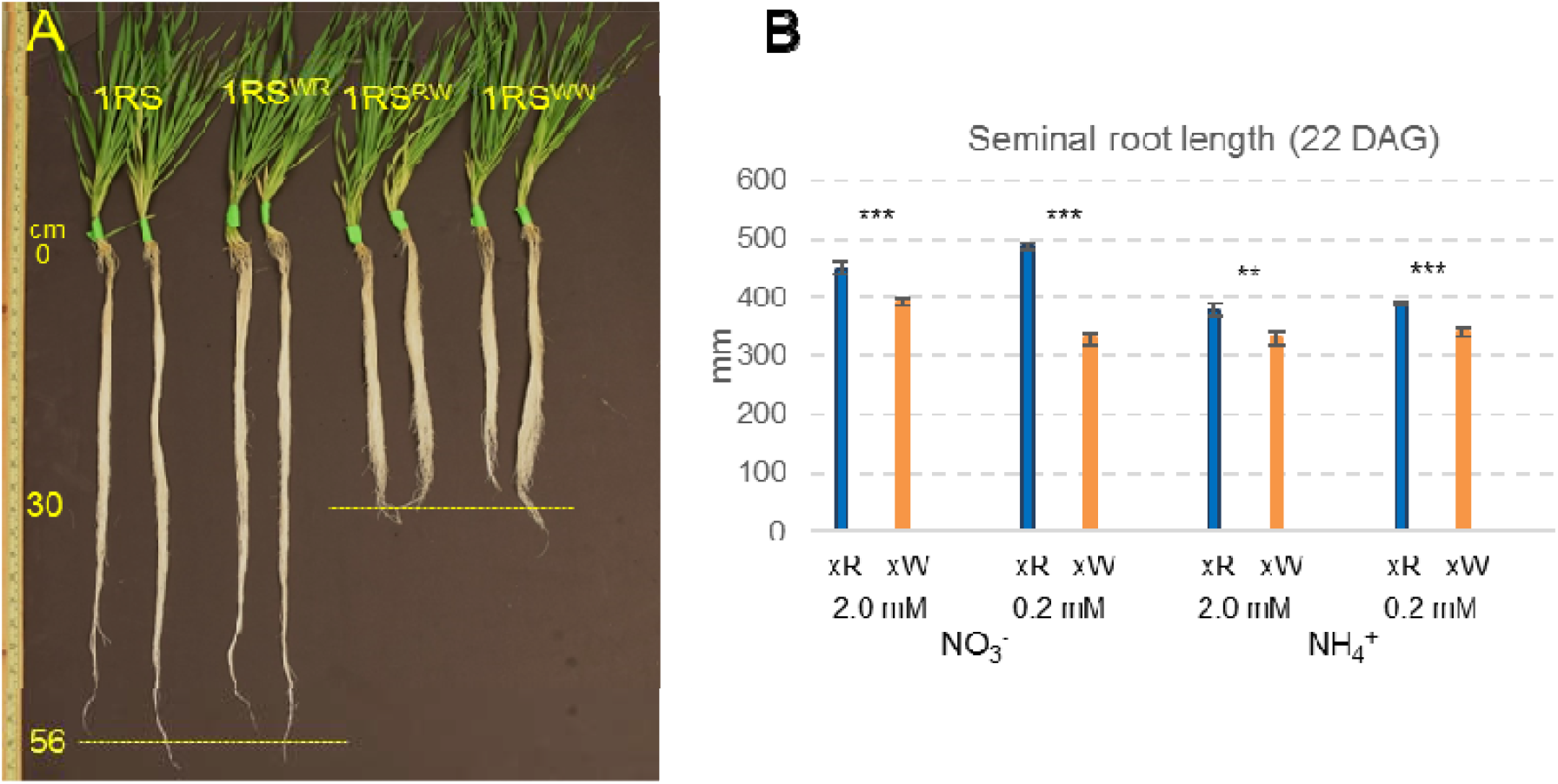
Differences in primary root length between plants carrying the distal rye (1RS and 1RS^WR^) and wheat chromosome segments (1RS^RW^ and 1RS^WW^) in hydroponic culture. **A**) Representative plants in normal culture medium. **B)** Seminal root length in hydroponic media using nitrate or ammonium at two different concentrations (2.0 mM and 0.2 mM) with n= 12 plants per genotype in each treatment (1RS^xR^ *vs.* 1RS^xW^).

#### Root elongation time course

The previous results indicated that the differences in primary root length between the 1RS^xR^ and 1RS^xW^ plants start at an early stage of root development. These differences were consistent between the lines with the same distal chromosome segment, so the following experiments were carried out using only the 1RS and 1RS^RW^ lines. To characterize the growth response in more detail, we performed daily measurements of roots and calculated elongation rates. Within each experiment, data was analyzed as repeated measures (split plot in time with conservative degrees of freedom). The experiment was repeated three times and overall ANOVAs were performed using experiments as a blocking factor, with replications nested within experiments (Supplementary Tables S16-S19).

In the overall ANOVAs, the two genotypes showed highly significant differences over time *(P* < 0.0001) in accumulated root length (Supplementary Table S16, Fig. 4A) and elongation rate (Supplementary Table S18, Fig. 4B). The differences between days and the interaction genotype by day interactions were also highly significant for both parameters (P < 0.001, Supplementary Tables S16 and S18). In separate ANOVAs performed for individual days, we detected significant differences in accumulated root length starting at 11 DAG, although differences were already close to significant at 10 DAG (P = 0.053, Supplementary Table S17, Fig. 4A). In the same experiments, the differences in the rate of elongation were significant from day 8.5 (P= 0.0328, Supplementary Table S19, Fig. 4B).

**Fig. 4.**
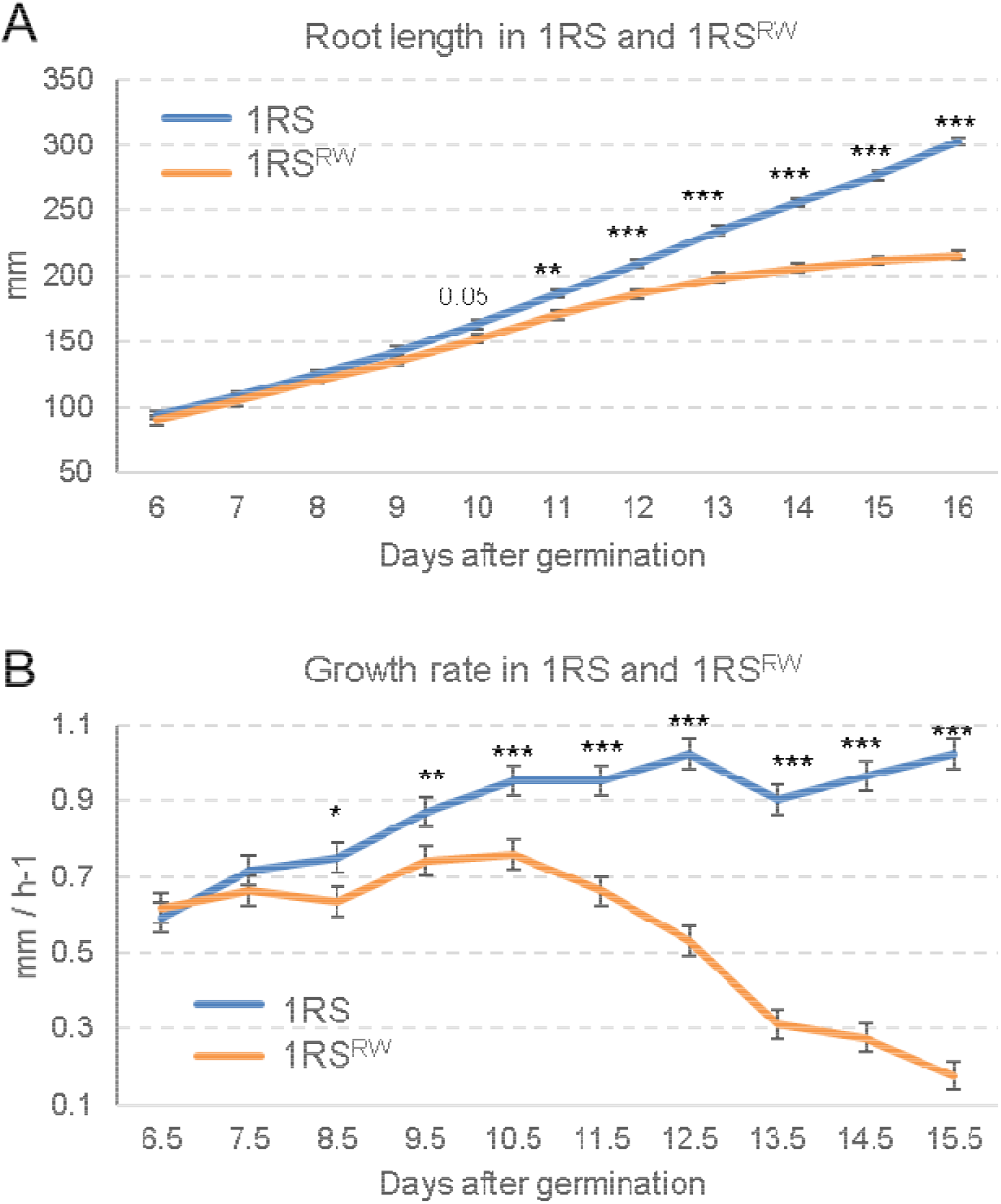
Time course of primary root growth of plants carrying the distal rye (1RS) or wheat chromosome segments (1RS^RW^) in hydroponics culture. LS means and standard errors are from the ANOVA of the three combined experiments. **A**) Cumulative length of the second longest seminal root (mm, Tables S16 and S17). **B**) Elongation rate of the second longest seminal root (mm/ h, Tables S18 and S19).* = *P* <0.05, ** = *P*<0.01, *** = *P*<0.001.

### 1RS^xR^ and 1RS^xW^ lines differ in root apical dominance and in the distribution of reactive oxygen species (ROS)

To test if the arrest in seminal root growth in the 1RS^RW^ plants was associated with a loss of seminal root apical dominance, we characterized the distribution of lateral roots. Figure 5 showed clear differences between the seminal roots of 1RS and 1RS^RW^ plants imaged at 22 DAG. In the seminal roots of the 1RS^xR^ plants, lateral roots started to appear after the first 90.1 ± 2.7 mm from the root tip suggesting active apical dominance (Fig. 5 and Supplementary Fig. S5). By contrast, the seminal roots of the 1RS^xW^ plants showed lateral roots appearing on average 14.7 ± 2.5 mm from the root apex. The differences between genotypes were highly significant (P < 0.0001, Supplementary Fig. S5).

**Fig. 5.**
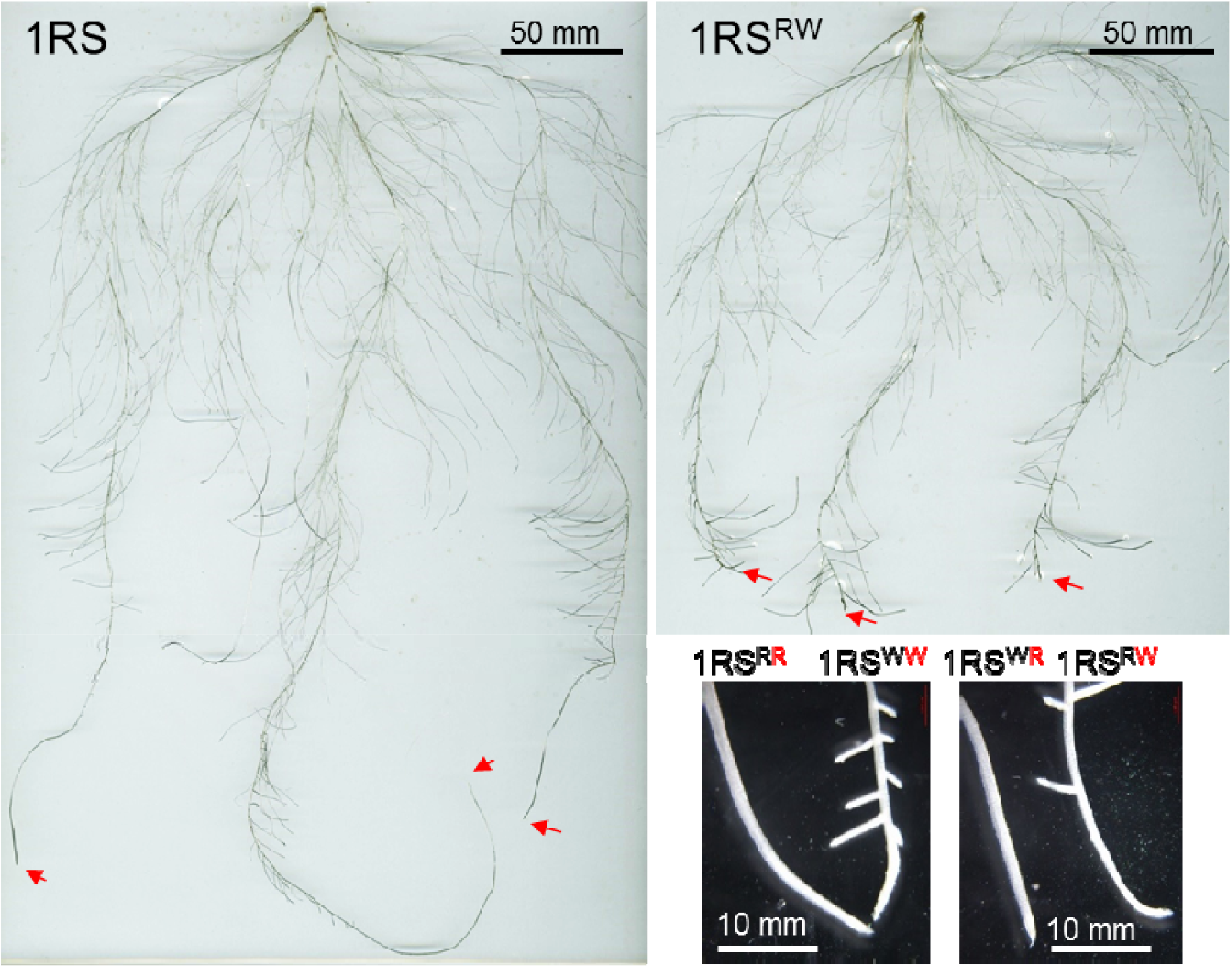
Development of laterals on the seminal roots of plants carrying the distal rye or wheat segments. Plants were grown for 22 days in hydroponics under low nitrogen conditions (0.2 mM nitrate). Arrows indicate the root apical meristem (RAM) of the seminal roots. The inset shows a detailed view of the seminal root tips for the four genotypes. 1RS^xW^ lines show lateral roots close to the RAM.

Since ROS gradients affect root elongation (Tyburski *et al*., 2009), we estimated their distribution in seminal roots 17 DAG by measuring the amount of formazan produced from the reduction of nitro blue tetrazolium (NBT) and fluorescence in DCF-DA staining. Formazan intensity, associated with superoxide anions, was similar in the distal region of the roots but was significantly lower in the 1RS^RW^ than in the 1RS roots proximal to 650 μm (*P* < 0.05), and even lower after 860 μm (*P* < 0.001, Fig. 6A). Fluorescence intensity from the DCF-DA staining, associated mainly with hydrogen peroxide, peroxynitrite and hydroxyl radicals, showed a very different pattern, with significantly higher intensities in 1RS^RW^ relative to the 1RS roots between 250 and 950 μm, and even higher differences between 350 and 640 μm (*P* < 0.001, Fig. 6B).

**Fig. 6.**
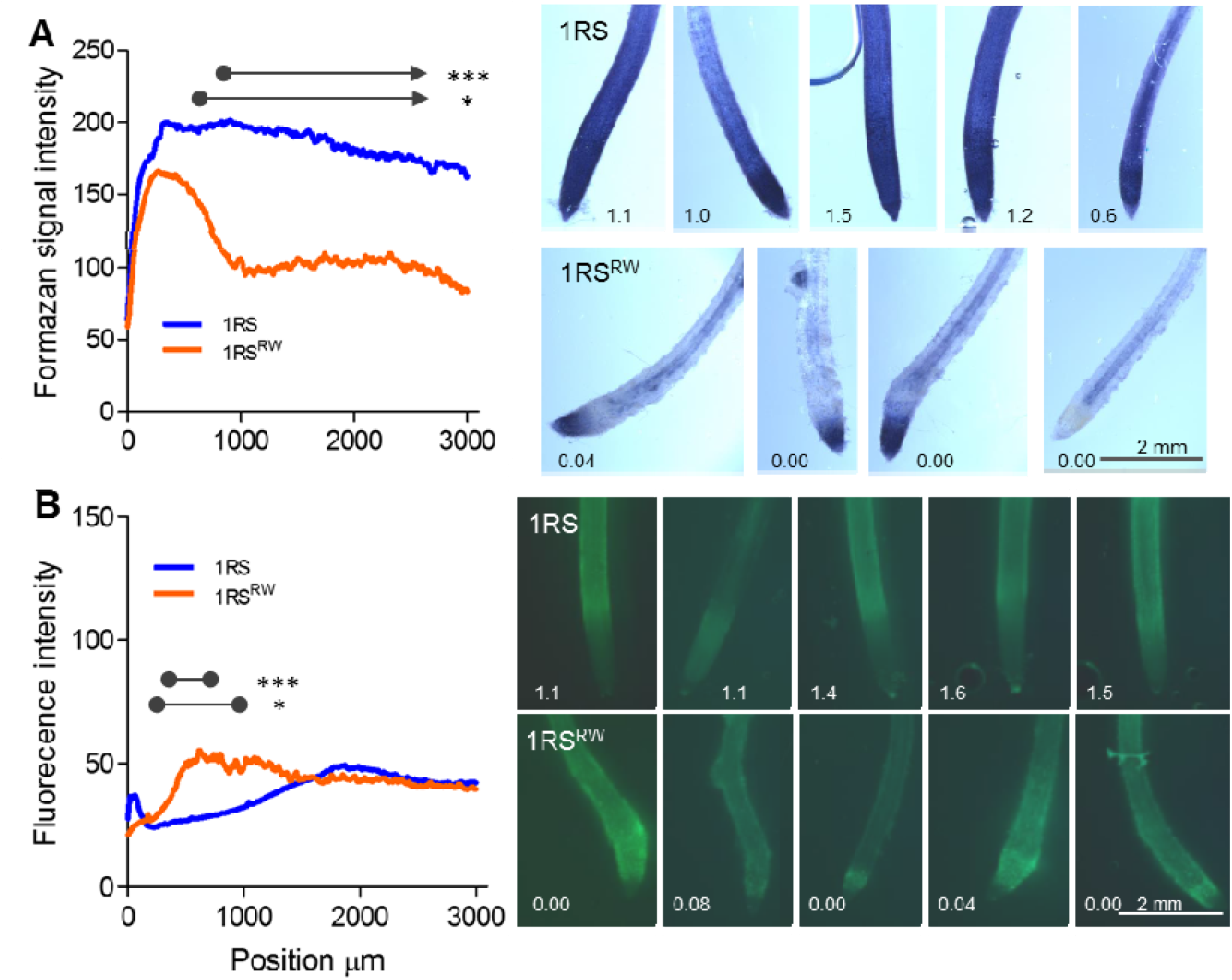
Fig. 6. Distribution of ROS along the main roots of plants carrying the distal rye (1RS) or wheat chromosome segments (1RS^RW^) 17 days after germination. A) Formazan signal intensity along the roots as determined with NBT (mainly detects superoxide anions). B) Fluorescence signal intensity along the root as determined by DCF-DA (mainly detects hydrogen peroxide). Representative NBT and DCF-DA stained roots from the corresponding experiment (number indicates growth rate in mm h^-1^ of the individual roots at the last sampling time). *= P < 0.05, ***= P < 0.001, n=5.

Finally, we compared the distribution of formazan during lateral root development in the same two genotypes (Supplementary Fig. S6). No differences between genotypes were detected at 6 DAG, but at 9 DAG NBT staining revealed lateral root primordia close to the RAM in the 1RS^RW^ but not in the 1RS line. By 15 DAG large lateral roots were formed close to the RAM in 1RS^RW^, but no lateral root primordia were observed in the same region in the 1RS line. These results confirmed the differences in apical dominance between the two genotypes.

## DISCUSSION

### 1RS^xR^ genotypes show deeper roots in the field and increased tolerance to waterlogging and terminal drought

Previous studies have shown an association between the introgression of the rye 1RS arm in wheat and improved resistance to water stress (Ehdaie *et al.*, 2012; Ehdaie *et al.*, 2003; Hoffmann, 2008; Moreno-Sevilla *et al.*, 1995; Zarco-Hernandez *et al.*, 2005). In three of these studies, the 1RS.1BL lines showed increased root biomass compared to the non-1RS control lines in large pot or sand-tube experiments. However, these differences were not validated in the field. In this study we showed that differences in grain yield and biomass between plants carrying a complete 1RS translocation and NILs with an introgressed distal wheat chromosome segment (1BS) are associated with differences in total root length density and average root diameter in the field. Field excavations across three blocks of a field experiment including the four different 1RS NILs provided an opportunity to visualize the differences in their root systems and to quantify these differences using horizontal soil cores at consistent depths.

This experiment confirmed the hypothesis that the 1RS^xR^ lines have a higher root density throughout the soil profile, with roots that reach deeper in the soil than the 1RS^xW^ lines (Fig. 2A). The more extensive root system provides a simple explanation for the higher carbon isotope discrimination and increased stomatal conductance values observed in the 1RS^xR^ plants in previous field experiments (Howell *et al.*, 2014). Through their deeper root system, the 1RS^xR^ plants can access more stored soil moisture, keep their stomata open longer, generate additional photosynthetic products and biomass than the 1RS^xW^ plants. This was reflected in this study in significantly higher NDVI values during the vegetative phase and increased grain yield values in the1RS^xR^ plants relative to the 1RS^xW^ plants, particularly under limiting irrigation (Fig. 1).

A previous study using eleven triticales showed that lines with better drought tolerance displayed root traits less affected by drought and waterlogging than the susceptible lines (Grzesiak *et al.*, 2002). In our study we observed that the 1RS^xR^ lines performed better than the 1RS^xW^ lines under both drought and waterlogging conditions (Fig. 1). The better performance of the 1RS^xR^ lines under waterlogging may be associated with their more extensive and deeper root system, but we cannot rule out the possibility that anatomical and/or physiological differences known to impact tolerance to waterlogging (Bailey-Serres and Voesenek, 2008) also affected the differences observed in this study. In wheat, short-term periods of waterlogging are known to affect the functionality of the root system (Malik *et al.*, 2001). Interestingly, an early study comparing one triticales and two wheat accessions after two cycles of waterlogging and drainage (seven days long each) found that the seminal roots of the triticale were twice as long as the seminal roots of the wheat cultivars (Thompson *et al.*, 1992). In a later study preformed in a stagnant medium that resulted in the death of the complete seminal root system, it was shown that the same triticale accession had a higher ratio of nodal roots/shoot fresh weight and a larger proportion of aerenchyma in root cross-section (18% vs. 14 and 12% in the two wheat cultivars). This trait was likely associated with improved O_2_ diffusion from shoots (Watkin *et al.*, 1998). Although the number of accessions included in the previous studies was too small to make any generalization, it indicates that additional studies will be required to determine if the 1RS^xR^ and 1RS^xW^ lines differ in anatomical or physiological traits that can affect their differential tolerance to waterlogging.

### 1RS^xW^ shows an earlier arrest of seminal root growth than 1RS^xR^ in hydroponic culture

The differences in root depth observed between the Hahn 1RS^xR^ and 1RS^xW^ NILs in the field were paralleled by drastic changes in seminal root length in hydroponic cultures (Fig. 3A). These differences were robust across experiments and were detected with different nitrogen sources and concentrations (Fig. 3B). We hypothesize that these early differences in seminal root length may have contributed to the observed differences in total root length density observed in the deepest soil core samples in the field (Fig. 2, Fig. 4B).

The early and consistent differences in root growth under controlled conditions, provided the opportunity to study the process in detail. During the first week of development, root growth occurred at the same rate for both genotypes, suggesting that the differences were not primarily associated with embryonically determined differences in root elongation. Instead, differences in root growth consistently manifested during the second week across multiple hydroponic experiments. The growth rate of the seminal roots of the 1RS^xW^ plants gradually decreased during the second and third week, to come close to zero by the end of the third week, whereas growth continued in the 1RS^xR^ plants (Fig. 4A). The consistent timing of these events suggests that these changes are developmentally regulated.

The growth arrest of the seminal roots in the 1RS^xW^ plants was accompanied by the proliferation of lateral roots in close proximity to the RAM, suggesting a loss of apical dominance. The RAM consists of a quiescent center (QC) surrounded by stem cells that generate new daughter cells, which undergo additional divisions in the proximal region of the meristem and differentiate in the transition zone (Heyman *et al.*, 2014). At a cellular level, a balance between cell proliferation and cell elongation/differentiation determines root growth rate (Pacifici *et al.*, 2015). The arrest of the growth of seminal roots in 1RS^xW^ plants suggests a modification cell proliferation and/or cell elongation/differentiation. Additional studies will be required to determine if this arrest involves changes in the QC and/or modifications in the root regions adjacent to the meristem. In any case, the dramatic reduction in growth and apical dominance argues for an early developmental program switch in the regulation of the RAM in the 1RS^xW^ plants.

### The distribution of ROS along seminal roots differed between 1RS^xW^ and 1RS^xR^ lines

The transition from cell proliferation to cell elongation and differentiation and the subsequent development of lateral roots depends on the distribution of ROS along the root axis, specifically on the opposing gradients of superoxide and hydrogen peroxide. Superoxide is predominant in dividing cells in the meristematic zone, while hydrogen peroxide is predominant in elongated cells in the differentiation zone (Dunand *et al.*, 2007; Tsukagoshi *et al.*, 2010; Voothuluru and Sharp, 2013; Yamada *et al.*, 2018). The balance between these reactive oxygen species modulates the transition between root proliferation and differentiation zones.

Seventeen days after germination, the apical region of the seminal roots of the 1RS lines showed opposing gradients of superoxide and hydrogen peroxide characteristic of elongating roots (Fig. 6). A different ROS distribution was detected in the arrested 1RS^xW^ roots, where superoxide was restricted to the distal ~700 μm and increased levels of DCF-DA fluorescence were detected between 250-950 μm in the cell proliferation zone (Fig. 6). The contrasting patterns of ROS distribution reflect the major developmental changes that differentiate the seminal roots of the 1RS and 1RS^xW^ genotypes.

Changes in ROS distribution can be triggered by the altered expression of major genes that control the size of the meristematic zone (De Tullio *et al.*, 2010). These genes include *UPBEAT1 (UPB1),* a basic helix-loop-helix (bHLH) transcription factor that regulates the meristematic zone size by restricting H_2_O_2_ distribution in the elongation zone (Tsukagoshi *et al.*, 2010). In addition, *ROOT MERISTEM GROWTH FACTOR 1 (RGF1)* and the transcription factor *RGF1 INDUCIBLE TRANSCRIPTION FACTOR 1 (RITF1)* that mediates *RGF1* signaling can modulate the distribution of ROS along the root developmental zones leading to enhanced stability of PLETHORA2 (PLT2)(Yamada *et al.*, 2018). Reduced expression of *PLETHORA* in the root apical region (Chen *et al.*, 2011) or changes in its distribution (Ercoli *et al.*, 2018) have been associated with impaired root growth.

It remains unknown if the differential pattern of ROS distribution in the roots of the 1RS^xW^ plants is the result of changes in the wheat homologs of these central developmental genes or a more direct effect on genes affecting the redox balance in different developmental root zones. The differences in superoxide and hydrogen peroxide distribution between the seminal roots of the 1RS^RW^ and 1RS plants were measured after the arrest in root growth (Fig. 6). Therefore, we currently do not know if the changes in ROS distribution are a cause or consequence of the changes observed in root growth and apical dominance.

### Conclusions and future directions

Results presented here indicate that the differences in grain yield between the 1RS^xW^ and 1RS^xR^ lines were preceded by significant differences in aerial biomass. These differences were generally larger under restricted or excessive irrigation than under normal irrigation, suggesting that the introgression of the distal wheat segment into the 1RS chromosome resulted in a reduced tolerance to these stresses.

The hydroponic studies showed that the introgression of the distal wheat chromosome segment in 1RS^xW^ resulted in drastic changes in apical dominance, location of lateral roots and ROS distribution along the roots. Since the 1RS^xW^ and 1RS^xR^ lines are highly isogenic (Howell *et al.*, 2014), they provide a useful tool to identify the genes that regulate this major developmental change in the wheat root apical meristem. The identification of these genes can not only contribute to our basic understanding of the regulation of root development in grasses, but also provide tools to modulate wheat root architecture and mitigate negative impacts of water stress in wheat production.

## Supporting information

Howell et al. Supplementary Figures and Tables

## ACKNOWLEDGEMENTS

This project was supported by USA-Israel BARD grant UC-4916-16, the Agriculture and Food Research Initiative Competitive Grant 2017-67007-25939 (WheatCAP) from the USDA National Institute of Food and Agriculture, by the International Wheat Yield Partnership (IWYP), and the Howard Hughes Medical Institute; and in Argentina by the CONICET and the ANPCYT (PICT 2014-1887).

## Supplementary information

### Supplementary Figures

Supplementary Fig. S1. Excavation of the first block including the four genotypes.

Supplementary Fig. S2. Plant biomass and grain yield.

Supplementary Fig. S3. Percent decrease in total root length density and diameter.

Supplementary Fig. S4. Differences between 1RS^xR^ and 1RS^xW^ in different root parameters.

Supplementary Fig. S5. Distance between the primary root apex and the first lateral root.

Supplementary Fig. S6. Distribution of nitro blue tetrazolium (NBT) staining during root development.

### Supplementary Tables

Supplementary Table S1. NDVI waterlogging 2013-2014.

Supplementary Table S2. NDVI waterlogging 2013-2014 by day.

Supplementary Table S3. Yield waterlogging 2013-2014.

Supplementary Table S4. NDVI waterlogging 2015-2016.

Supplementary Table S5. NDVI waterlogging 2015-2016 by day.

Supplementary Table S6. Yield waterlogging 2015-2016.

Supplementary Table S7. NDVI drought 2015-2016.

Supplementary Table S8. NDVI drought 2015-2016 by day.

Supplementary Table S9. Yield drought 2015-2016.

Supplementary Table S10. Total root length density.

Supplementary Table S11. Total root length density contrast.

Supplementary Table S12. Total root length density contrast by depth.

Supplementary Table S13. Average root diameter.

Supplementary Table S14. Average root diameter contrast.

Supplementary Table S15. Average root diameter contrast by depth.

Supplementary Table S16. Repeated measures for root length Supplementary Table S17. Root length (mm) by day.

Supplementary Table S18. Repeated measures for root elongation rate.

Supplementary Table S19. Root elongation rate by day.

