## Supplementary material for "A wheat/rye polymorphism affects seminal root length and is associated with drought and waterlogging tolerance": Howell et al. Supplementary Figures and Tables

**SUPPLEMENTARY FILE**

**Supplementary Figures**

**Supplementary Fig. S1. Excavation of the first block including the four genotypes** (1RS, 1RS^RW^, 1RS^WR^, 1RS^WW^). The holes below the center of each plot (20 to 140 cm deep) indicate were the horizontal soil core samples were taken. Samples from 160 cm and 180 cm were taken from the other two blocks after we discovered the presence of roots at 140 cm in block 1. Plants were at the tillering stage at the time of the root sampling.

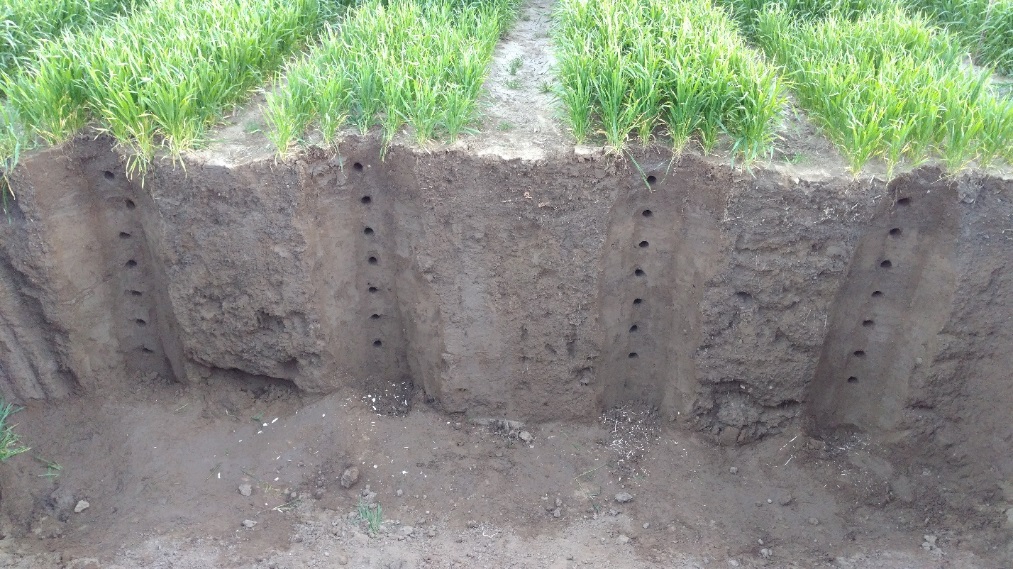

20 cm

40 cm

60 cm

80 cm

100 cm

120 cm

140 cm

**1RS	1RS^RW^	1RS^WR^	1RS^WW^**

**Supplementary Fig. S2. Plant biomass and grain yield.** Plant biomass estimated by NDVI (A) and grain yield (B) under normal irrigation (control), waterlogging and drought conditions. Plants with the distal rye segment include the original Hahn-1RS lines (RR) and the line with the proximal wheat segment (WR =1RS^WR^). Plants with the distal wheat segment include the line with both wheat segments (WW= 1RS^WW^) and with only the distal wheat segment (RW =1RS^RW^). Note the similar response of the lines with a similar distal chromosome segment. Differences between genotypes were generally larger under water stress conditions.

A

2016 NDVI drought

|  | RR | RW | WR | WW |
| --- | --- | --- | --- | --- |
| Control | 0.921746 | 0.853352 | 0.921405 | 0.870643 |
| Drought | 0.903559 | 0.807638 | 0.898808 | 0.796361 |

2016 NDVI waterlogging

|  | RR | RW | WR | WW |
| --- | --- | --- | --- | --- |
| Control | 0.921746 | 0.853352 | 0.921405 | 0.870643 |
| Flood | 0.904608 | 0.816504 | 0.895001 | 0.823806 |

2014 NDVI waterlogging

|  | RR | RW | WR | WW |
| --- | --- | --- | --- | --- |
| Control | 0.860153 | 0.783159 | 0.867762 | 0.799751 |
| Flood | 0.816502 | 0.694181 | 0.813326 | 0.702762 |

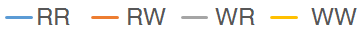

B

2014 Yield drought

|  | RR | RW | WR | WW |
| --- | --- | --- | --- | --- |
| Control | 7952.07 | 5678.27 | 7325.9 | 6204.77 |
| Flood | 6868.6 | 5114.23 | 6907.2 | 4980.13 |

2016 Yield waterlogging

|  | RR | RW | WR | WW |
| --- | --- | --- | --- | --- |
| Control | 7952.07 | 5678.27 | 7325.9 | 6204.77 |
| Flood | 8430.27 | 5480 | 7804.9 | 5411.6 |

2014 Yield waterlogging

kg/ / ha

|  | RR | RW | WR | WW |
| --- | --- | --- | --- | --- |
| Control | 6750 | 4454 | 6519 | 4906.5 |
| Flood | 6360.5 | 3462.5 | 6027 | 3539.5 |

**Supplementary Fig. S3. Percent difference in total root length density and diameter.** Differences between genotypes expressed as a percent of the 1RS^xR^ values (((1RS^xR^ **-** 1RS^xW^) / 1RS^xR^) *100). **A**) Percent difference in total root length. **B**) Percent difference in average root diameter. Error bars represent standard errors of the means across blocks.

A

B

% decrease in total root length density (difference relative to 1RSxR)

| Depth (cm) | % difference in total length |
| --- | --- |
| 40 | 55.4383% |
| 60 | 41.4068% |
| 80 | 39.1732% |
| 100 | 63.8965% |
| 120 | 74.263% |
| 140 | 93.1077% |
| 160 | 96.5475% |
| 180 | 100% |

% descrease in root diameter (diffference relative to 1RSxR)

% decrease in diameter

| Depth (cm) |  |  |
| --- | --- | --- |
| 40 | 4000% | -11.1003% |
| 60 | 6000% | 8.35525% |
| 80 | 8000% | 11.4426% |
| 100 | 10000% | -5.97262% |
| 120 | 12000% | 74.263% |
| 140 | 14000% | 89.3473% |
| 160 | 16000% | 81.6932% |
| 180 | 18000% | 100% |

**Supplementary Fig. S4. Differences between 1RS^xR^ and 1RS^xW^ in different root parameters with depth**. The experiment is organized in a split plot design with 4 genotypes as main plots and depths as subplots. **A**) Root surface (SQRT (x+0.5)). **B**) Root volume ((x+1)^-4^). **C**) Number of root tips (SQRT (x+0.5)). **D**) Number of root forks (SQRT (x+0.5)). *P* values were calculated using contrasts between 1RS^xR^ and 1RS^xW^ lines in individual ANOVAs by depth. Error bars represent standard errors of the means across blocks. *P* values are from transformed data (to meet the assumptions of the ANOVA) and means and SE are untransformed.

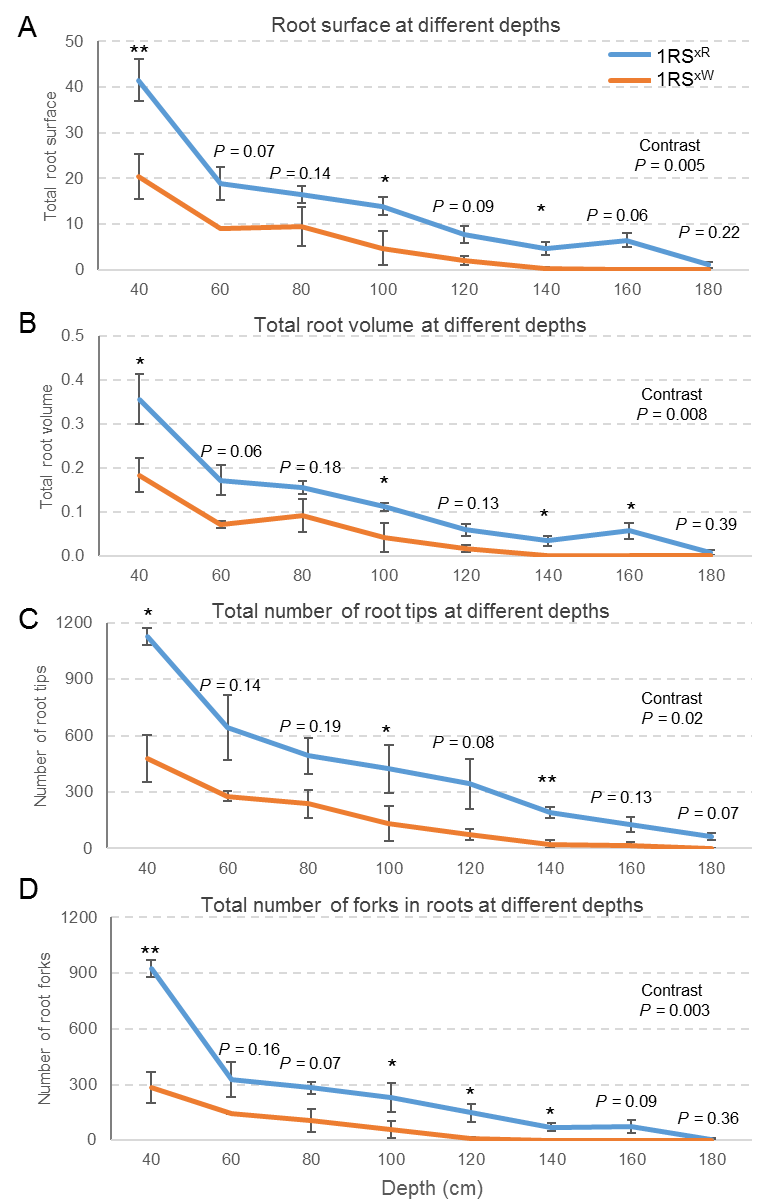

**Supplementary Fig. S5. Distance between the primary root apex and the first lateral root** in lines carrying the distal rye (1RS and 1RS^WR^) or wheat segment (1RS^WW^ and 1RS^RW^) in plants grown in hydroponics conditions for 22 days after germination. The lateral roots appeared significantly closer to the primary root apical meristem in the lines with the distal wheat segment than in the lines with the distal rye segment (*P =* 2.2 E-15, total n=23).

1RS	1RS^RW^	1RS^WR^	1RS^WW^

Distance between RAM and lateral roots

|  | mm |
| --- | --- |
| 1RS | 88.4 |
| 1RSRW | 11.4 |
| 1RSWR | 91.8 |
| 1RSWW | 18.7 |

**Supplementary Fig. S6.** Distribution of nitro blue tetrazolium (NBT) staining during root development in plants grown under low nitrate conditions (0.2 mM nitrate). No differences were detected between 1RS and 1RS^RW^ lines six days after germination (DAG). At 9 DAG, stained primordia of the lateral roots close to the root apical meristem (RAM) are evident in 1RS^RW^ but not in the 1RS. By 15 DAG, lateral roots are formed close to the RAM in 1RS^RW^ but no lateral root initiation is observed in the 1RS line close to the RAM.

NBT staining

1RS	1RS^RW^

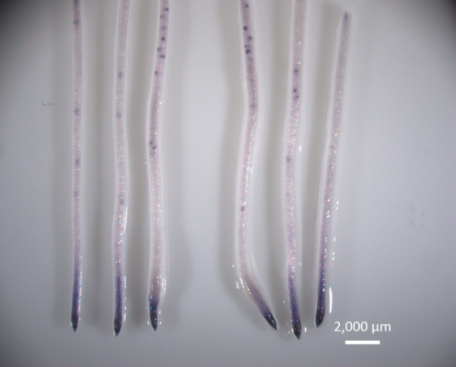

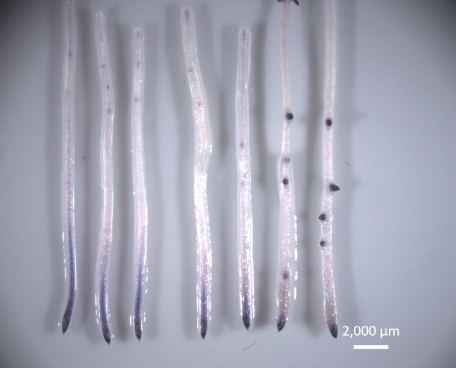

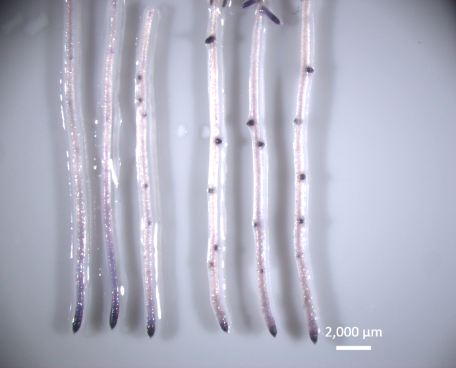

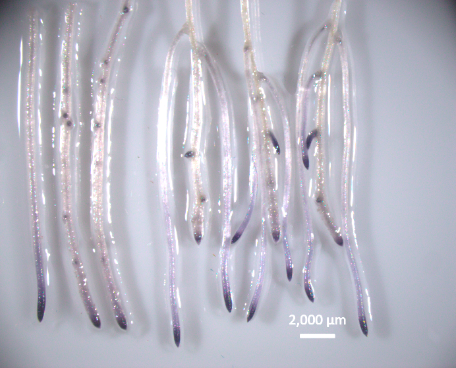

**6 d**

**9 d**

**12 d**

**15 d**

**Supplementary Tables**

**Supplementary Table S1a. NDVI waterlogging 2013-2014.** Split plot RCBD repeated in time ANOVA table for the Normalized Difference Vegetation Index (NDVI). Main plots are waterlogging and normal irrigation, subplots are the genotypes 1RS^xR^ (averaged 1RS and 1RS^WR^) and 1RS^xW^ (averaged 1RS^RW^ and 1RS^WW^), and sub-sub plots are days. Since there is only one degree of freedom for days, we did not calculated conservative degrees of freedom.

```
Source                    dF     Type III SS   Mean Sq.      F Value    Pr > F
Main plot
Treat                      1       0.04034      0.04034       90.30     0.0025
Block                      3       0.00447      0.00149        3.34     0.1743
Error: Block*Treat         3       0.00134      0.00045

Genot.                     1       0.07140      0.07140      407.17     <.0001
Treat*Genot                1       0.00386      0.00386       22.02     0.0034
Error: Block*Treat*Genot.  6       0.00105      0.00018

day                        1       0.13770      0.13770     1130.10     <.0001
day*Treat                  1       0.00100      0.00100        8.25     0.0232
day*Genot                  1       0.00618      0.00618       50.75     <.0001
day*Treat*Genot            1       0.00007      0.00007        0.60     0.4518
Error                     12       0.00146      0.00012
R-square= 0.99456
```

**Supplementary Table S2**. **NDVI waterlogging 2013-2014 by day**. Because the day * genotype was significant we also analyzed the individual split plot ANOVAs by individual days. The significance of treatment and block effects was tested using the Block*Treatment as error (df = 3) and the significance of Genotype and Treatment * Genotype using the residual error (df = 6)

| Source | df | April 17  *P* | April 30  *P* |
| --- | --- | --- | --- |
| Treat | 1 | <.0001 | 0.0083 |
| Block | 3 | 0.0126 | 0.3453 |
| Genotype | 1 | <.0001 | <.0001 |
| Treat*Genot | 1 | 0.0032 | 0.0097 |

**Supplementary Table S3. Yield waterlogging 2013-2014.** Split plot RCBD ANOVA table for grain yield. 2013-2014 field experiments at Davis, CA. Main plots are waterlogging and normal irrigation and subplots are the genotypes 1RS^xR^ (averaged 1RS and 1RS^WR^) and 1RS^xW^ (averaged 1RS^RW^ and 1RS^WW^).

```
Source                   dF     Type III SS    Mean Sq.    F Value    Pr > F
Treat                     1       2623671      2623671       3.56    0.1556
Block                     3        510586       170195       0.23    0.8701
Block*Treat               3       2210786       736929

Genot                     1      21593447     21593447     138.98    <.0001
Treat*Genot               1        545493       545493       3.51    0.1101
Error                     6        932244       155374
R-square= 0.967193
```

**Supplementary Table S4.** **NDVI waterlogging 2015-2016.** Split plot RCBD repeated in time ANOVA table for the Normalized Difference Vegetation Index (NDVI). Main plots are waterlogging and normal irrigation, subplots are genotypes with 1RS^xR^ (averaged 1RS and 1RS^WR^) and 1RS^xW^ (averaged 1RS^RW^ and 1RS^WW^), and sub-sub plots are the four days. Since it is not possible to randomize days, *P* values were calculated using conservative degrees of freedom (indicated in blue, and calculated by dividing by df days= 3).

```
Source                     df   Type III SS   Mean Sq.    F Value     Pr > F
Main plot
Treat                       1     0.01214     0.01214     960.33      0.0010
Block                       2     0.00307     0.00154     121.57      0.0082
Error: Block*Treat          2     0.00002     0.00001

Genot                       1     0.05815     0.05815     266.34      <.0001
Treat*Genot                 1     0.00121     0.00121       5.53      0.0783
Error: Block*treat*Genot.   4     0.00087     0.00022

                     df  Con.df                                              P Con.df
day                   3     1     0.14303     0.14303     894.6     <.0001    <.0001
day*Treat             3     1     0.00161     0.00161      10.1     0.0002    0.0130
day*Genot             3     1     0.02108     0.02108     131.5     <.0001    <.0001
day*Treat*Genot       3     1     0.00059     0.00059       3.7     0.0253    0.0906
Error                24     8     0.00128     0.00005
R-square= 0.9947
```

**Supplementary Table S5. NDVI waterlogging 2015-2016 by day**. Because the day * genotype was significant we also analyzed the individual split plot ANOVAs by individual days. The significance of treatment and block effects was calculated using the Block*Treatment as error (df = 2) and the significance of Genotype and Treatment * Genotype using the residual error (df = 4). The four NDVI measurement were taken before the breakage of the irrigation pipeline, so are not affected by this accident.

| Source | df | March 24  *P* | April 6  *P* | April 13  *P* | April 28  *P* |
| --- | --- | --- | --- | --- | --- |
| Treat | 1 | 0.0110 | 0.0057 | 0.0133 | 0.0146 |
| Block | 2 | 0.0383 | 0.0291 | 0.2118 | 0.0972 |
| Genotype | 1 | 0.0080 | 0.0008 | 0.0009 | <.0001 |
| Treat*Genot | 1 | 0.9137 | 0.0595 | 0.0687 | 0.1587 |

**Supplementary Table S6.** **Yield waterlogging 2015-2016.** Split plot RCBD ANOVA table for grain yield. Main plots are waterlogging and normal irrigation, whereas subplots are genotypes with 1RS^xR^ (averaged 1RS and 1RS^WR^) or 1RS^xW^ (averaged 1RS^RW^ and 1RS^WW^). The broken irrigation pipeline increased variability and contributed to the absence of treatment differences.

```
Source                     df     Type III SS    Mean Sq.    F Value    Pr > F
Treat                       1           218          218       0.00    0.9772
Block                       2       4048744      2024372       9.58    0.945
Block*Treat                 2        422468       211234       0.44    0.6702
Genot                       1      14317650     14317650      30.03    0.0054
Treat*Genot                 1        711945       711945       1.49    0.2888
Error                       4       1906973       476743
R-square= 0.91092
```

**Supplementary Table S7.** **NDVI drought 2015-2016.** Split plot RCBD repeated in time ANOVA table for the Normalized Difference Vegetation Index (NDVI). Main plots are terminal drought and normal irrigation, subplots are the genotypes 1RS^xR^ (averaged 1RS and 1RS^WR^) and 1RS^xW^ (averaged 1RS^RW^ and 1RS^WW^), and sub-sub plots are the four days. Since it is not possible to randomize days, P values were calculated using conservative degrees of freedom (in blue, calculated by dividing by df days= 3).

```
Source                    df     Type III SS   Mean Sq.   F Value   Pr > F
Main plot
Treat                      1      0.019388    0.019388      7.38     0.1130
Block                      2      0.000617    0.000309      0.12     0.8949
Error: Block*Treat         2      0.005254    0.002627

Genot.                     1      0.075617    0.075617    171.84     0.0002
Treat*Genot                1      0.004705    0.004705     10.69     0.0308
Error: Block*treat*Genot.  4      0.001760    0.000440

                     df  Con.df                                             P Con.df
day                   3     1     0.231431    0.077144    434.07     <.0001  <.0001
day*Treat             3     1     0.017311    0.005770     32.47     <.0001  0.0005
day*Genot             3     1     0.025883    0.008628     48.55     <.0001  0.0001
day*Treat*Genot       3     1     0.000529    0.000176      0.99     0.4134    0.35
Error                24     8     0.004265    0.000178
R-square= 0.98897
```

**Supplementary Table S8. NDVI drought 2015-2016 by day**. Because the day * genotype was significant we also analyzed the individual split plot ANOVAs by individual days. The significance of treatment and block effects was calculated using the Block*Treatment as error (df = 2) and the significance of Genotype and Treatment * Genotype using the residual error (df = 4)

| Source | df | March 24  *P* | April 6  *P* | April 13  *P* | April 28  *P* |
| --- | --- | --- | --- | --- | --- |
| Treat | 1 | 0.2372 | 0.3538 | 0.2148 | 0.0562 |
| Block | 2 | 0.8203 | 0.7723 | 0.8138 | 0.7577 |
| Genotype | 1 | 0.0128 | 0.0049 | 0.0011 | <.0001 |
| Treat*Genot | 1 | 0.0984 | 0.1885 | 0.1559 | 0.0005 |

**Supplementary Table S9.** **Yield drought 2015-2016.** Split plot RCBD ANOVA table for grain yield. Main plots are waterlogging and normal irrigation and subplots are the genotypes 1RS^xR^ (average 1RS and 1RS^WR^) and 1RS^xW^ (average 1RS^RW^ and 1RS^WW^).

```
Source                     df    Type III SS   Mean Sq.    F Value    Pr > F
Treat                      1       2030506     2030506	0.83	   0.4587
Block                      2        961895      480948	0.20	   0.8359
Block*treat                2       4900514     2450257

Genot                      1       9389144     9389144     22.81 	   0.0088
Treat*Genot                1         15394       15394      0.04	   0.8561
Error                      4       1646190      411547
R-square= 0.9131
```

**Supplementary Table S10. Total root length density**. Split plot – RCBD ANOVA for the excavation experiment including genotypes as main plots (averages of 1RS and 1RS^WR^ and 1RS^RW^ and 1RS^WW^) and depths as subplots. Conservative degrees of freedom (in blue) were used to test the significance of Depth and Genotype * Depth because depths cannot be randomized. This approach is very conservative and the correct *P* value is between the two estimates. This does not affect the significance of the differences between genotypes, which is the main objective of this analysis.

```
Source                    df    Type III SS   Mean Sq.  F Value   Pr > F
Main plot
Genot                      1      64176       64176     36.58     0.0263
Block                      2      13794        6897      3.93     0.2028
Main error Block*Genot     2       3509        1754

Subplots             df Cons.df                                         P Cons.df
Depth                 7    1     327458       46780      25.07    <.0001   0.007
Genot*Depth           7    1      31705        4529       2.43    0.0497   0.190
Subplot Error        24   3.4     44781        1866
R-square= 0.806
```

**Supplementary Table S11.** **Total root length density contrast**. The previous analysis (Table S10) has limited degrees of freedom for genotype (ν_1_= 1ν_2_= 2), so we performed an alternative ANOVA including the four genotypes and a statistical contrast between the lines with distal rye and wheat segments (ν_1_= 1,ν_2_= 6).

```
Source                    df     Type III SS   Mean Sq.     F Value    Pr > F
Contrast distal R vs. W    1       504.23       504.23      19.64      0.004
Error Block*Gen            6       154.07        25.68
```

^1^ Data transformed (SQRT(length +0.05) to optimize homogeneity of variances and normality of residuals.

**Supplementary Table S12. Total root length density contrast by depth**. Alternative ANOVA including the four genotypes and a statistical contrast between the lines with distal rye and wheat segments by depth.

| Source | 40 cm | 60 cm | 80 cm | 100 cm | 120 cm | 140 cm | 160 cm | 180 cm |
| --- | --- | --- | --- | --- | --- | --- | --- | --- |
| Contrasts distal R *vs.* W | 0.0020 | 0.1279 | 0.1313 | 0.0347 | 0.0844 | 0.0180 | 0.0678 | 0.1687 |

^1^ Data transformed (SQRT(length +0.05) to optimize homogeneity of variances and normality of residuals.

**Supplementary Table S13. Average root diameter**. Split plot RCBD ANOVA table for the excavation experiment using genotypes as main plots (averages of 1RS and 1RS^WR^ and 1RS^RW^ and 1RS^WW^) and depths as subplots. Conservative degrees of freedom (in blue) were used to test the significance of Depth and Genotype * Depth. This is a very conservative approach and the correct *P* value is between the two estimates. This does not affect the significance of the differences between genotypes, which is the main objective of this analysis.

```
Source                 df        Type III SS   Mean Sq.    F Value    Pr > F
Main plot
Genot                   1          0.0083442   0.0041721    13.92    0.0649
Block                   2          0.1096073   0.1096072     0.53    0.6536
Main error Block*Genot  2          0.0157427   0.0078713

Subplots                df cons.df                                         p cons.df
Depth                   7     1    0.3666487   0.0523784    19.45   <0.0001   0.02
Gen*Depth               7     1    0.1232096   0.0176014     6.54    0.0002   0.08
Subplot Error          24    3.4   0.0646355   0.0026931
R-square= 0.68817
```

**Supplementary Table S14.** **Average root diameter contrast.** Alternative split plot ANOVA including the four genotypes and a statistical contrast between the lines with distal rye and wheat segments (ν_1_= 1ν_2_= 6). More sensitive than Table S13.

```
Source                       df     Type III SS   Mean Sq.    F Value    Pr > F
Contrast distal R vs. W      1       0.21715       0.21715     16.70      0.006
Main error Block*Genot       6       0.07800       0.01300
```

**Supplementary Table 15. Average root diameter contrast by depth.** Alternative ANOVA including the four genotypes and a statistical contrast between the lines with distal rye and wheat segments by depth.

| Source | 40 cm | 60 cm | 80 cm | 100 cm | 120 cm | 140 cm | 160 cm | 180 cm |
| --- | --- | --- | --- | --- | --- | --- | --- | --- |
| Contrasts distal R *vs.* W | 0.3596 | 0.7262 | 0.5726 | 0.8369 | 0.0735 | 0.0012 | 0.0256 | 0.0287 |

**Supplementary Table 16. Repeated measures for root length (mm)**. ANOVA between 1RS and 1RS^RW^ of the second longest main root measured daily between 6 and 16 d. Within each experiment, data was analyzed as a split plot with genotype as main plot and days as subplot (using conservative df, indicated in blue). The three experiments were combined in a single analysis using experiment as block and reps nested in experiment (Fig. 4A).

```
Source              df         Type III SS   Mean Sq.    F Value    Pr > F
Exp                  2           143301.9    71650.9      22.35     <.0001
genot                1            97054.5    97054.5      30.27     <.0001
genot*rep(Exp)      47           150674.8     3205.8

                    df Cons.df                                            P cons.df
day                 10     1     1688462     168846    1225.49     <.0001   <.0001
genot*day           10     1      101971      10197      74.01     <.0001   <.0001
Error              490    49       67512      137.8
R-square= 0.96986
```

**Supplementary Table S17. Root length (mm) by day**. ANOVAs for each time point combining the three experiments (used as blocks). Untransformed LS means (n=10) obtained from ANOVAs by day.

|  | **6** | **7** | **8** | **9** | **10** | **11** | **12** | **13** | **14** | **15** | **16** |
| --- | --- | --- | --- | --- | --- | --- | --- | --- | --- | --- | --- |
| **1RS** | 90.9 | 104.8 | 122.2 | 139.5 | 160.2 | 183.5 | 206.3 | 231.9 | 253.0 | 274.3 | 299.6 |
| 1RSRW | 88.6 | 102.8 | 118.8 | 133.3 | 150.9 | 169.3 | 184.9 | 197.9 | 204.9 | 210.6 | 214.6 |
| ANOVA *P* | 0.446 | 0.581 | 0.420 | 0.173 | 0.053 | 0.006 | 0.0002 | <.0001 | <.0001 | <.0001 | <.0001 |

**Supplementary Table S18. Repeated measures for root elongation rate (mm/ h^-1^)**. ANOVA between 1RS and 1RS^RW^ for the second longest main root measured daily between 6 and 16 d. Within each experiment, data was analyzed as a split plot with genotype as main plot and days as subplot (using conservative df, indicated in blue). The three experiments were combined in a single analysis using experiment as block and reps nested in experiment (Fig. 4B).

```
Source                    df     Type III SS    Mean Sq.  F Value    Pr > F
Exp                        2        3.3099      1.6550       8.95    0.0005
genot                      1       22.9752     22.9752     124.23    <.0001
genot*rep(Exp)            47        8.6921      0.1849
                       Cons.df                                              P Cons.df
day                  9     1        5.9958      0.6662      13.18    <.0001  0.0007
genot*day            9     1       13.5174      1.5019      29.72    <.0001  <.0001
Error              441    49       22.2881      0.0505
R-square=0.7109
Data was transformed to the power of **1.7 to improve normality of residuals
```

**Supplementary Table S19. Root elongation rate by day.** ANOVAs for each time point combining the three experiments (used as blocks). Untransformed LS means (n=10) obtained from ANOVAs by day.

|  | **6.5** | **7.5** | **8.5** | **9.5** | **10.5** | **11.5** | **12.5** | **13.5** | **14.5** | **15.5** |
| --- | --- | --- | --- | --- | --- | --- | --- | --- | --- | --- |
| **1RS** | 0.5794 | 0.7116 | 0.7418 | 0.8622 | 0.9449 | 0.9441 | 1.0136 | 0.8946 | 0.9556 | 1.0128 |
| 1RS^RW^ | 0.5997 | 0.6545 | 0.6245 | 0.7335 | 0.7487 | 0.6455 | 0.5141 | 0.2973 | 0.2602 | 0.1630 |
| ANOVA *P* | 0.7085 | 0.2923 | 0.0328 | 0.0021 | 0.0006 | <.0001 | <.0001 | <.0001 | <.0001 | <.0001 |

P values from transformed data to the power of **1.7 to improve normality of residuals.
